# Identification of Bovine Genotypes Conferring Diminished Susceptibility to Salmonellosis and Colonization by *Salmonella* and *E. coli* O157:H7

**DOI:** 10.1101/2020.07.06.190751

**Authors:** Kristi L. Anderson, Duane K. Ramsey, Tim A. Day, Steve A. Carlson

## Abstract

*Salmonella* and *E. coli* O157:H7 are two of the most important problems for the beef industry. Cattle can develop salmonellosis and persistently harbor *Salmonella*, or they can asymptomatically shed *Salmonella* and/or *E. coli* O157:H7 resulting in contamination of the hide and carcass surfaces during processing. Additionally, *Salmonella* infiltrates lymph nodes that get incorporated into ground beef. In this study, we investigated the possibility of identifying cattle with reduced susceptibility to one or both of these infections. Empirical observations from previous studies suggested that a diminished susceptibility was possible in amelanotic cattle, *i.e*., cattle bearing the *mcr1/mcr1* genotype and lacking overt black pigmentation. By searching for single nucleotide polymorphisms (SNP) present in the 34 genes encoding the *Salmonella* interactome, we identified a SNP that was consistently present in amelanotic cattle with diminished susceptibility to *Salmonella*. Specifically, we used an *ex vivo* assay to screen 500 cattle blood samples for the diminished ability of *Salmonella* to penetrate peripheral leukocytes. Diminished *Salmonella* penetration was observed in 150 of these blood samples and 147 of these samples harbored two alleles bearing a SNP that introduces a miRNA cleavage site (*bta-let-7b*) in the 3’UTR of the *bsynJ1* gene, which we designate as the *SYNJ1/SNYJ1* genotype. Further *ex vivo* studies revealed a decreased expression of *SYNJ1* in leukocytes bearing the *SYNJ1/SNYJ1* genotype. *In vivo* experimental challenge studies revealed a diminished susceptibility to salmonellosis in cattle with the *SYNJ1/SNYJ1::mcr1/mcr1* genotype. Additional *in vivo* challenge studies revealed that *SYNJ1/SNYJ1::mcr1/mcr1* cattle have a decreased susceptibility to lymph node infiltration by two *Salmonella* serotypes (*S*. Anatum and *S*. Montevideo) implicated in this lymph node problem, and a decreased susceptibility to *E. coli* O157:H7 colonization of the recto-anal junction. A field study revealed that the *SYNJ1/SNYJ1::mcr1/mcr1* genotype was five times more prevalent, when compared to the *SYNJ1/synj1::mcr1/mcr1* and *synj1/ synj1::mcr1/mcr1* genotypes, in *Salmonella*-free lymph nodes. Small-scale genetic surveys revealed that the *SYNJ1/SNYJ1* genotype was present in the following *mcr1/mcr1* breeds: Akaushi, Barzona, Braunvieh, Hereford, Piedmontese, Red and White Holsteins, Red Angus, Red Poll, Shorthorn, Simmental (Red), and Tarentaise. Studies using the aforementioned *ex vivo* penetration assay, which putatively predicts the diminished susceptibility phenotype, revealed that the penetrance of the diminished susceptibility is >99% in *SYNJ1/SNYJ1::mcr1/mcr1* cattle but only ∼1% in *SYNJ1/SNYJ1* cattle with at least one *MCR1* allele. Further studies with the *ex vivo* assay revealed that three additional SNPs are part of a genotype conferring diminished susceptibility to a broad array of *Salmonella* serotypes commonly associated with cattle. In summary, the studies presented herein reveal a bovine genotype associated with decreased susceptibility to *Salmonella* and *E. coli* O157:H7. PSR Genetics LLC holds a U.S. patent on testing for the *SYNJ1/SNYJ1* genotype (patent number 9,049,848) while the three complementary SNPs are under further investigation.

## 1. Introduction

In cattle, *Salmonella* occasionally cause clinical disease but this pathogen can also be asymptomatically harbored in the intestinal tracts and lymph nodes of cattle. The latter phenomenon is a food safety concern since lymph nodes serve as a protective conduit for *Salmonella* passage into ground beef (Brichta-Harhay et al., 2008). The bovine recto-anal junction is a depot for *E. coli* O157:H7, which is a commensal in cattle but highly pathogenic in humans (Smith et al., 2014).

Given the need to mitigate these two food safety problems and the animal health problems associated with salmonellosis, identifying novel intervention strategies is critical and the aim of this study is to identify cattle that are less susceptible to infections by these two bacteria. Twelve *in vivo* experimental *Salmonella* challenge studies (Brewer et al., 2014; Carlson et al., 2007; Carlson et al., 2002a; Carlson et al., 2002b; Carlson et al., 2005; McCuddin et al., 2008; Rasmussen et al., 2005; Wu et al., 2002; Xiong et al., 2013; Xiong et al., 2012; Xiong et al., 2011; Xiong et al., 2010) revealed that certain calves were less likely to be successfully infected with *Salmonella* in an experimental setting. Across these 12 studies there were 13 calves that were excluded from the studies since these animals could not be successfully colonized or did not elicit salmonellosis after experimental challenge. All 13 calves were from the following amelanotic breeds: Hereford, Jersey, Red Angus, and Red and White Holstein.

The core recessive genotype of amelanotic breeds is *mcr1/mcr1* that leads to a non-functional or non-expressed melanocortin 1 receptor (Seo et al., 2007). The intact and fully functional version of the receptor activates a tyrosinase that converts phaeomelanin to eumelanin that provides black pigmentation. In the absence of functional melanocortin 1 receptors, phaeomelanin predominates thus yielding a yellowish/red pigmentation (Seo et al., 2007). Other alleles, such as dilution alleles in white breeds like Charolais (Gutiérrez-Gil et al., 2007), provide other non-black color patterns in *mcr1/mcr1* cattle (Seo et al., 2007).

Since we were able to empirically associate the *mcr1/mcr1* genotype with a putative phenotype for diminished susceptibility to *Salmonella*, the aims of this study were to search for single nucleotide polymorphisms (SNP) present in the 34 genes encoding proteins exploited by *Salmonella* during the infection process, *i.e*., the *Salmonella* interactome (Schleker et al., 2012b). We first used an *ex vivo* peripheral leukocyte *Salmonella* penetration assay to find leukocytes that “resisted” *Salmonella* penetration and then we looked for single nucleotide polymorphisms (SNPs) present in the 34 genes encoding the *Salmonella* interactome. Once a SNP was correlated with the *ex vivo* “resistance” phenotype, we obtained calves with zero, one, or two copies of the SNP and performed two different *in vivo* experimental *Salmonella* challenges. One experimental challenge involved a serotype (*S*. Newport) that causes overt salmonellosis in calves while the other challenge involved two serovars (*S*. anatum and *S*. Montevideo) implicated in the lymph node infiltration problem (Brichta-Harhay et al., 2012), along with a co-challenge of *E. coli* O157:H7. A field study was also performed to investigate the presence of the SNP in cattle lymph nodes that were free of *Salmonella*.

Small-scale genetic screening was also performed in order to uncover the prevalence of the SNP in various populations of cattle. Studies with samples from melanotic cattle (*MCR1/MCR1* or *MCR1/mcr1*) were also performed in order to compare the penetrances of the diminished susceptibility in amelanotic and melanotic cattle. Herein we report the discovery of a SNP that is present in *mcr1/mcr1* cattle with diminished susceptibility to *Salmonella* and *E. coli* O157:H7. This diminished susceptibility to *Salmonella* extends to a broad array of *Salmonella* serotypes if three additional SNPs are present. PSR Genetics LLC has a proprietary claim on the SNP present in the *Salmonella* interactome (US patent number 9,049,848) while the three complementary SNPs are under further investigation.

## 2. Materials and Methods

### 2.1. Ex vivo screening for decreased susceptibility to Salmonella

Since previous 12 *in vivo* studies suggested that amelanotic cattle could be a population to look for animals with decreased susceptibility to *Salmonella*, we obtained blood samples from approximately 500 amelanotic cattle from the following breeds or crosses of these breeds: Braunvieh, Hereford, Piedmontese, Red & White Holsteins, Red Angus, Shorthorn, Simmental, and Tarentaise. As a Control, blood was taken from about the same number of melanotic cattle representing the Angus and Black & White Holstein breeds. About 5mL of whole blood was collected from the caudal vein, placed in EDTA tubes and immediately used in the *ex vivo Salmonella* susceptibility assay. Blood was subjected to centrifugation and the erythrocyte fraction was removed. Approximately 10^7^ colony-forming units (**CFUs**) of a standard *S. enterica* serotype Typhimurium strain [**SL1344**; (Wray and Sojka, 1978)] was then added to the cells. As a Control, cells were incubated with non-invasive standard *S. enterica* serotype Typhimurium strain BJ68 (Penheiter et al., 1997). After 12 hours of incubation at 37°C, extracellular (non-invasive) bacteria were killed by adding an equivalent volume of PBS containing 100Μg/ml (final concentration equal to 50Μg/ml) of gentamicin-a bactericidal antibiotic that rapidly kills extracellular bacteria but does not penetrate eukaryotic cells (Gianella et al., 1973). Samples were then incubated at 37°C for 1 hour to ensure that all non-invasive bacteria are killed. Leukocytes were centrifuged and the gentamicin-containing media was removed and replaced with 50ΜL of phosphate-buffered saline containing 1% Triton which lyses the leukocytes. Lysates were then plated on XLD agar that was incubated overnight at 37°C. The following day, black-centered colonies were enumerated and the invasion/survival of *Salmonella* was then calculated as a percent that equals 100(number of *Salmonella* recovered from within leukocytes/10^7^).

### 2.2. Screening for SNPs in the 34 genes encoding the Salmonella interactome

A literature search revealed the existence of a SNP in the 3’ UTR of the gene encoding synaptojanin (Cargill et al., 2008), a protein that *Salmonella* exploits during the invasion process (Marcus et al., 2001). To screen for this SNP that is a C to T substitution at nucleotide 3981, we used a PCR assay (Forward oligonucleotide = 5’-AACCACCAGAGTAACAGACTACAC-3’; Reverse oligonucleotide = 5’-ATGCAGCTTACAGAACTCAGAGT-3’) to amplify a 362bp region encompassing the SNP. Dideoxynucleotide sequencing was used to evaluate PCR amplicons for the presence of the SNP, with an ambiguous signal indicating heterozygousity as per Fig. 2. We hereby designated the C to T substitution at nucleotide 3981 as the *SYNJ1* allele while the “wild-type” is the *synj1* allele.

**Figure 1.**
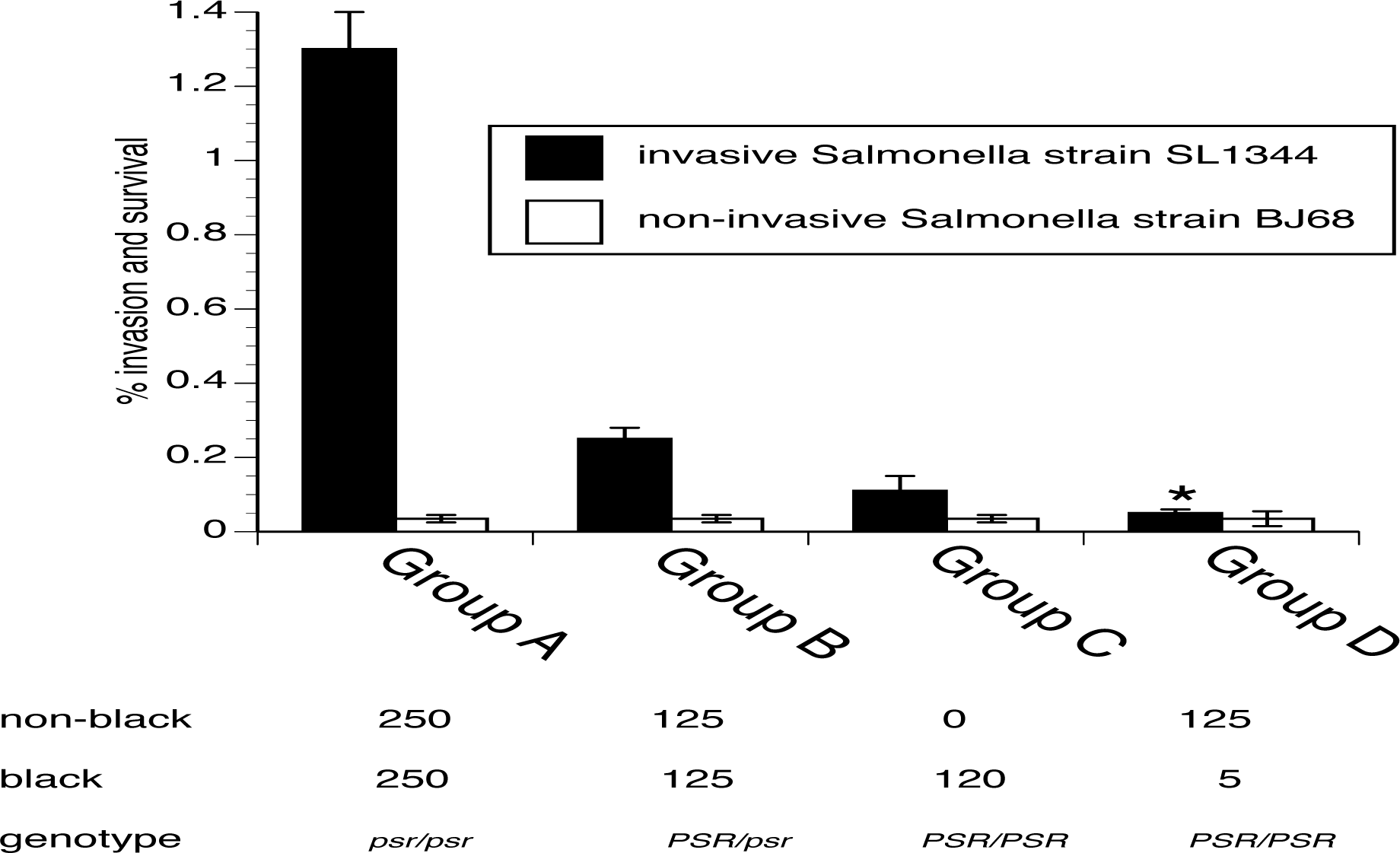
*Ex vivo* studies identifying amelanotic cattle blood cells that are less susceptible to infection by *Salmonella*. Data represent are the mean ± SEM for the % invasion and survival of *Salmonella* based on an arbitrary clustering of results. Numbers below the Groups represent the number of amelanotic and melanotic cattle represented in the given Group-specific data set. The bottom line of the embedded Table depicts the predominate synaptojanin genotype in each Group. \**P* > 0.05 versus the non-invasive strain.

**Figure 2.**
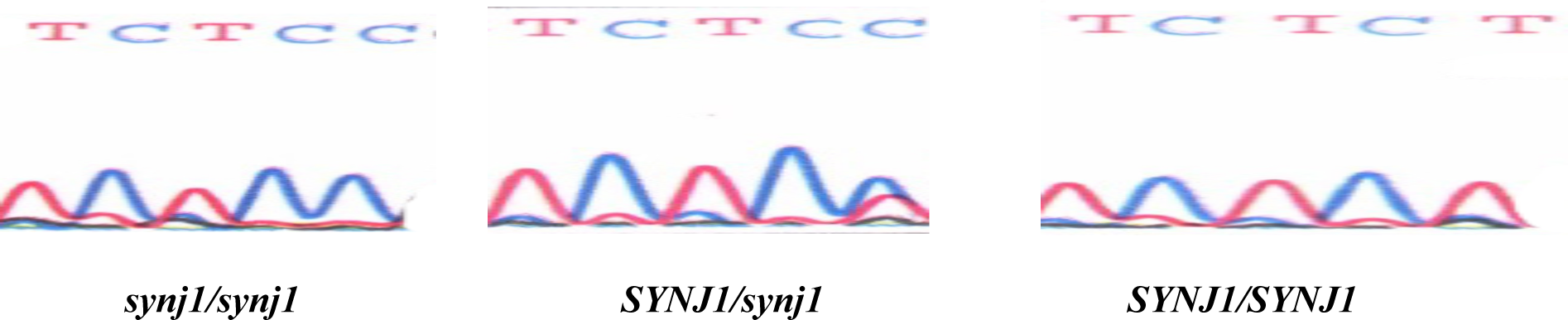
Chromatograms of the 3’UTR DNA sequencing obtained from cattle bearing the *synj1/synj1, SYNJ1/synj1, and SYNJ1/SYNJ1* genotypes. The region surrounding nucleotide 3981 (the far-right nucleotide in each pane) was PCR-amplified and then subjected to standard dideoxynucleotide sequencing. The terminal cytosine (C) is replaced with a thymidine (T) in one allele in the heterozygous sequence (both red AND blue peaks) and in both alleles in the *SYNJ1/SYNJ1* sequence.

### 2.3. Assessment of Synaptojanin Gene Expression in Leukocytes

Semi-quantitative RT-PCR was performed on blood samples obtained from cattle. RNA was isolated using the Blood RNEasy kit (Qiagen) and RT-PCR was performed as described previously described (Carlson et al., 2007). Our empirical studies revealed that amplification was consistently variable across the three genotypes between 10 and 15 cycles of PCR. As such, we used 12 cycles as an arbitrary time point to stop the reaction and visualize the amplicons using agarose gel electrophoresis.

### 2.4. In vivo experimental challenge with Salmonella Newport

Calves (2-6 months of age) with the nine possible combinations of *synj1, SYNJ1, MCR1*, and *mcr1* genotypes were obtained and orally challenged with 109 CFUs/kg of a multi-resistant strain of *Salmonella* Newport as per previous studies (Brewer et al., 2014; Carlson et al., 2007; Carlson et al., 2002a; Carlson et al., 2002b; Carlson et al., 2005; McCuddin et al., 2008; Rasmussen et al., 2005; Wu et al., 2002; Xiong et al., 2013; Xiong et al., 2012; Xiong et al., 2011; Xiong et al., 2010). Each day, calves were evaluated for four clinical signs of salmonellosis-*Salmonella* fecal shedding (assessed by qualitative fecal culture using XLD agar), diarrhea, pyrexia (rectal temperature ≥103.5°F), and ≥4% dehydration based on a skin tent test. An arbitrary clinical score was ascribed daily whereby a +1 was given for each of the four clinical parameters, *i.e*., a score equal to 4 indicates the simultaneous observation of all four signs on a given day. Animals were immediately euthanized (IM xylazine followed by IV pentobarbital) if any of the following were observed: recumbency, anorexia, profuse diarrhea, rectal temperature > 106oF, or dehydration >6%. Animals requiring euthanasia were ascribed a clinical score equal to 5 on the day of euthanasia. This experimental protocol was performed twice using three calves for each of the nine possible genotypes in each experiment, *i.e*., a total of 54 calves. For *SYNJ1/SYNJ1::mcr1/mcr1* cattle, a third experiment was performed with five calves that received a ten-fold higher dose (1010 CFUs/kg) of *S*. Newport.

### 2.5. In vivo experimental challenge with Salmonella Anatum and Montevideo

Calves (2-6 months of age) possessing one of the nine possible combinations of *synj1, SYNJ1, MCR1*, and *mcr1* genotypes were obtained and orally challenged with 109 CFUs/kg of a 1:1 cocktail of multi-resistant strains of *Salmonella* Anatum and Montevideo (Brichta-Harhay et al., 2012) as per previous studies (Brewer et al., 2014; Carlson et al., 2007; Carlson et al., 2002a; Carlson et al., 2002b; Carlson et al., 2005; McCuddin et al., 2008; Rasmussen et al., 2005; Wu et al., 2002; Xiong et al., 2013; Xiong et al., 2012; Xiong et al., 2011; Xiong et al., 2010). Each day, calves were evaluated for unexpected clinical signs of salmonellosis. On day 14 post-challenge, calves were euthanized (IM xylazine followed by IV pentobarbital) and six lymph nodes (superficial cervical, subiliac, popliteal, tuberal, gluteal, and ileocecal) were recovered from each calf. Lymph nodes were then subjected to quantitative culture of *Salmonella* as per Feye *et al*. (Feye et al., 2016). This experimental protocol was performed once using 12 calves with the *SYNJ1/SYNJ1::mcr1/mcr1* genotype and 16 calves representing the other eight possible genotype combinations of *synj1, SYNJ1, MCR1*, and *mcr1*.

### 2.6. Assessment of SNYJ1 Genotypes in Cattle with Salmonella-free Lymph Nodes

In order to correlate the absence of *Salmonella* with the *synj1, SYNJ1, MCR1*, and *mcr1* genotypes, *Salmonella*-free lymph nodes (n=200) from a Control group from a prior study (Feye et al., 2016) were subjected to PCR-based genotyping targeting the *synj1, SYNJ1, MCR1*, and *mcr1* alleles. DNA was extracted from the lymph nodes, and then subjected to the PCR-based sequencing procedure described herein.

### 2.7. In vivo experimental challenge with E. coli O157:H7

Calves experimentally infected with *S*. Anatum and Montevideo were also infected with 10^10^ CFUs/kg of *E. coli* O157:H7 strain isolated from our recent study (Feye et al., 2016). On day 14 post-challenge, approximately 0.3 gm of recto-anal junction scrapings were collected and transferred into enrichment broth (Sharma and Casey, 2014) and an aliquot of the broth was plated on sorbitol-MacConkey agar, incubated overnight at 37°C, and subjected to enumeration by manual counting of non-fermenting colonies the next day. From each genotype-specific set of agar plates, 96 colonies were selected and subjected to the PCR targeting *E. coli* O157:H7 virulence genes (Sharma and Casey, 2014). Load was then determined as (colonies recovered times the dilution factor times the percent of colonies yielding an *E. coli* O157H7-specific amplicon)/gm of feces. Prevalence was calculated as percent of harboring any *E. coli* O157:H7 within a genotype, and was compiled across genotypes.

### 2.8. Small-scale Assessment of synj1 and SYNJ1 Genotypes in Various Cattle Breeds

Blood was obtained from the following *mcr1/mcr1* breeds: Akaushi (n=24), Barzona (n=30), Braunvieh (n=100), Hereford (n=3), Piedmontese (n=35), Red and White Holsteins (n=20), Red Angus (n=160), Red Poll (n=40), Shorthorn (n=4), Red Simmental (n=20), and Tarentaise (n=40). Blood was also collected from the following *MCR1* breeds: Black and White Holsteins (n=250), Black Angus (n=250), and Black Simmental (n=5). The *ex vivo* invasion assay was performed using the blood samples in order to determine the penetrance of the diminished susceptibility relative to the presence of the *SYNJ1/SYNJ1* genotype.

### 2.9. Small-scale Assessment of Diminished Susceptibility to an Array of Salmonella serotypes

The *ex vivo* invasion/survival assay was performed using blood obtained from 100 *SYNJ1/SYNJ1::mcr1/mcr1* cattle whereby a broad array of *Salmonella* serotypes were used as the test strains. Specifically, we created and used a pool of 100 *Salmonella* laboratory strains representing 70 different serotypes known to be isolated from cattle based on a literature search. These serotypes include *S*. Dublin, *S*. Derby, *S*. Reading, *S*. Lubbock, *etc*. As a Control, the same pool of *Salmonella* were incubated with blood obtained from *synj1/synj1::MCR1/MCR1* cattle.

Since this pool of *Salmonella* were able to survive within approximately 50% of the blood samples obtained from *SYNJ1/SYNJ1::mcr1/mcr1* cattle, we examined potential genotypic differences between *SYNJ1/SYNJ1::mcr1/mcr1* cattle whose leukocytes “resisted” the pool of *Salmonella* serotypes when compared to *SYNJ1/SYNJ1::mcr1/mcr1* cattle whose leukocytes were most permissive of the invasion/survival of the *Salmonella* pool. To do so, blood samples were submitted to GeneSeek for 50k SNP analysis. Consensus SNPs were then obtained by comparing the SNP profiles of “hypo-susceptible” and “susceptible” animals using a proprietary algorithm.

### 2.10. Statistical Analyses

Statistical comparisons were made using an analysis of variance with Tukey’s *ad hoc* test for multiple comparisons (GraphPad Prism, Version 6, La Jolla, CA). Significant differences were defined at *P* ≤ 0.05.

## 3. Results

### 3.1. Ex vivo-based identification of cattle with decreased susceptibility to Salmonella

Since our previous 12 *in vivo* studies suggested that amelanotic cattle could be a population to look for animals with decreased susceptibility to *Salmonella*, we obtained blood samples from approximately 500 amelanotic cattle and 500 melanotic cattle. Blood was collected and used in an *ex vivo Salmonella* susceptibility assay that measures the eukaryotic cell invasion and intra-cellular survival of *Salmonella*. Interestingly, the results clustered into four different arbitrary groups representing a hierarchy of decreased susceptibility to *Salmonella*. As shown in Fig. 1, the maximal reduction in susceptibility to *Salmonella* was identified in some of the amelanotic cattle (Group D) while the vast majority of the melanotic cattle clustered into the most susceptible group (Group A). Blood from four melanotic cattle exhibited a moderate decrease in susceptibility (Groups B and C), while one melanotic animal yielded blood displaying the maximal resistance (Group D).

### 3.2. Screening for SNPs in the 34 genes encoding the Salmonella interactome

In order to correlate the results depicted in Fig. 1 with a genotype, we examined the literature for genetic variants in genes encoding the *Salmonella* interactome (Schleker et al., 2012b). A literature search revealed the existence of a SNP in the 3’ UTR of the gene encoding synaptojanin (Cargill et al., 2008), a protein that *Salmonella* exploits during the invasion process (Marcus et al., 2001). This SNP introduces part of an RNAi-cleavage site (*bta-let-7b*) into the gene, and this site could reduce the expression of this gene (Cargill et al., 2008). Using blood obtained from studies presented in Fig. 1, we used a PCR-based assay (Fig. 2) to identify the relative prevalence of this SNP (*i.e*., the *SYNJ1* allele) in the four susceptibility Groups. As shown in the bottom line of the Table embedded in Fig. 1, the *SYNJ1/SYNJ1* predominates Groups C and D. The heterozygous genotype (*SYNJ1/synj1)* predominates Group B while Group A is mostly *synj1/synj1*.

### 3.3. Assessment of Synaptojanin Gene Expression in Leukocytes

Since the studies presented in Fig. 1 revealed a possible association between decreased susceptibility and decreased expression of a bovine protein that *Salmonella* exploits during the infection process, we assessed the expression of the gene in leukocytes obtained from cattle exhibiting the various levels of susceptibility to *Salmonella*. RNA was isolated (and later Group-specifically pooled) from blood representing three of the Groups (Group C was not used because of the paucity of numbers). As shown in Fig. 3, synaptojanin gene expression was the lowest in Group D and the highest in Group A.

**Fig. 3.**
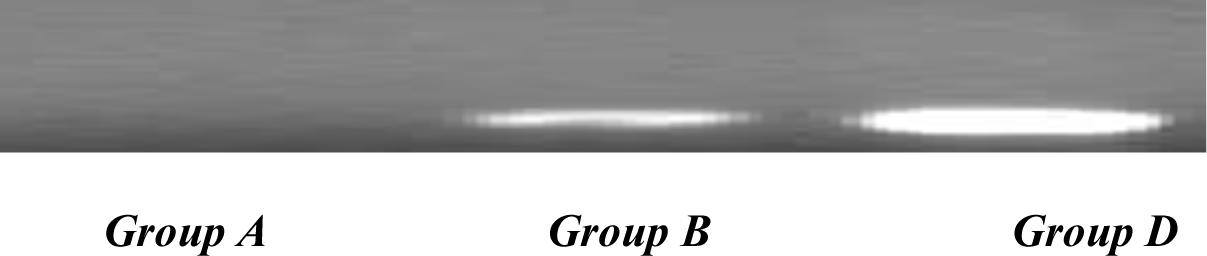
Semi-quantitative analysis of synaptojanin expression in blood samples obtained from cattle representing the susceptibility Groups A, B, and D that have the predominate genotypes *synj1/synj1, SYNJ1/synj1*, and *SYNJ1/SYNJ1*, respectively. RNA was isolated from blood and semi-quantitative RT-PCR was performed using 12 cycles of amplification to delineate differences in synaptojanin mRNA transcript levels.

### 3.4. Assessment of in vivo susceptibilities to Salmonella Newport across the SYNJ1 and synj1 genotypes

Since the preliminary *in vitro* and *ex vivo* studies revealed a possible link between decreased susceptibility to *Salmonella* in amelanotic calves possessing the *SYNJ1* allele, calves with the nine possible combinations of *SYNJ1, synj1, MCR1*, and *mcr1* genotypes were orally challenged with multi-resistant *Salmonella* Newport. Over the next two weeks, calves were then evaluated daily for four clinical signs of salmonellosis. As shown in Fig. 4, calves with the *SYNJ1/SYNJ1::mcr1/mcr1* genotype were least susceptible to salmonellosis. Specifically, euthanasia was required for all calves with any of the other eight genotypes whereas none of the *SYNJ1/SYNJ1::mcr1/mcr1* required euthanasia. Furthermore, increasing the dose 10-fold had no effect on the clinical outcome in *SYNJ1/SYNJ1::mcr1/mcr1* cattle.

**Fig. 4.**
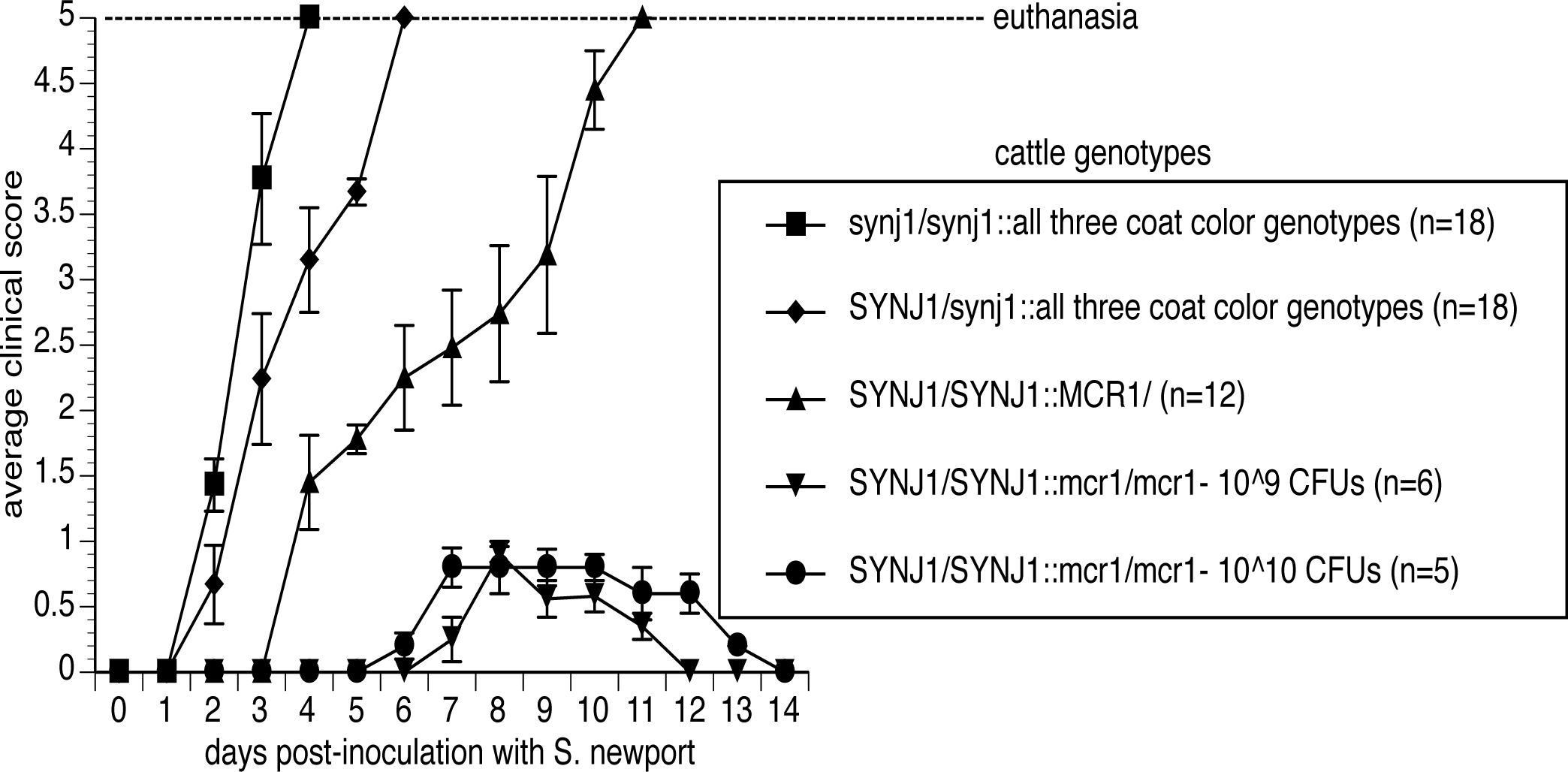
*Salmonella* Newport infectivity of cattle in accordance with the *SYNJ1* and *MCR1* genotypes. Two to six month-old calves were orally infected with 10^9^ (n=six animals/genotype) or 10^10^ (n=5 animals) CFUs of *Salmonella* Newport. Each day after infection on day0, calves were monitored for signs of salmonellosis-fecal shedding of *Salmonella*, diarrhea, fever, and anorexia. The clinical score is an accumulation of scores for these four parameters in which a +1 is ascribed for each parameter observed on a given day. Calves requiring euthanasia were ascribed a score of five.

### 3.5. Assessment of in vivo susceptibilities to Salmonella Anatum and Montevideo across the SYNJ1 and synj1 genotypes

Calves were orally challenged with *Salmonella* Anatum and Montevideo. On day 14 post-challenge, calves were euthanized and lymph nodes were recovered and subjected to quantitative culture of *Salmonella*. As shown in Fig. 5, *Salmonella* infiltration of the lymph nodes was markedly reduced in the *SYNJ1/SYNJ1::mcr1/mcr1* calves when compared to calves representing the other eight possible genotype combinations of *SYNJ1, synj1, MCR1*, and *mcr1*.

**Fig. 5.**
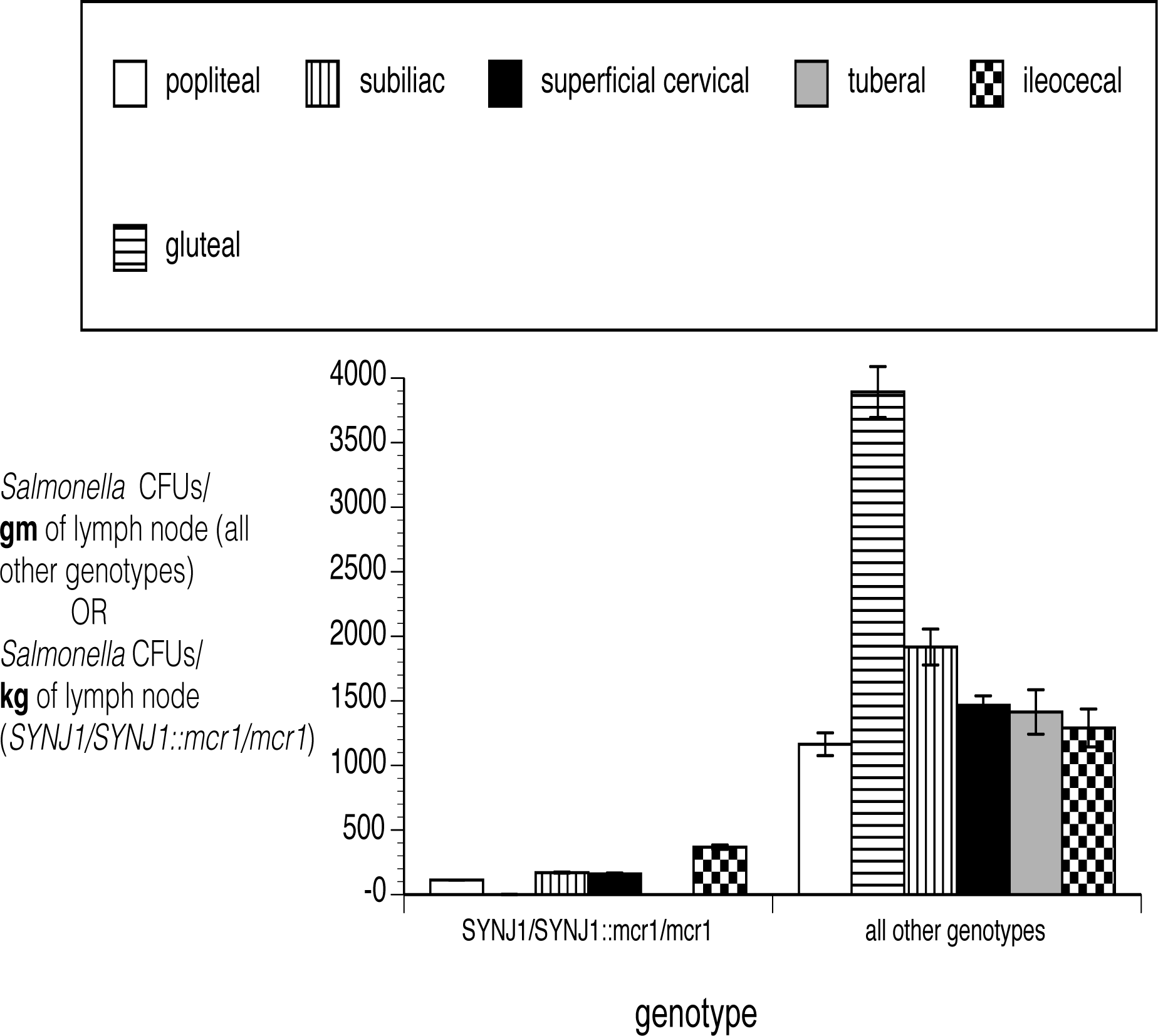
Assessment of lymph node infiltration of *Salmonella* in *SYNJ1/ SYNJ1::mcr1/mcr1* Cattle. Calves were experimentally infected with *Salmonella* Anatum and Montevideo and lymph nodes were recovered and *Salmonella* were enumerated in each lymph node. Data are presented, for scaling purposes, as *Salmonella* CFUs/gm or kg of lymph node for cattle of all other genotypes and *SYNJ1/SYNJ1::mcr1/mcr1* cattle, respectively. n=12 for the *SYNJ1/ SYNJ1::mcr1/mcr1* and n=16 for the other eight genotypes (*i.e*., two calves per genotype).

### 3.6. Assessment of snyj1 Genotypes in Cattle with Salmonella-free Lymph Nodes

In order to correlate the absence of *Salmonella* with the *SYNJ1, synj1, MCR1*, and *mcr1* genotypes, *Salmonella*-free lymph nodes from the Control group in a recent study (Feye et al., 2016) were subjected to PCR-based genotyping targeting the *SYNJ1* and *mcr1* genes. As shown in Fig. 6, the *Salmonella* load and prevalence were significantly lower in cattle with *SYNJ1/ SYNJ1::mcr1/mcr1* genotype.

**Figure 6.**
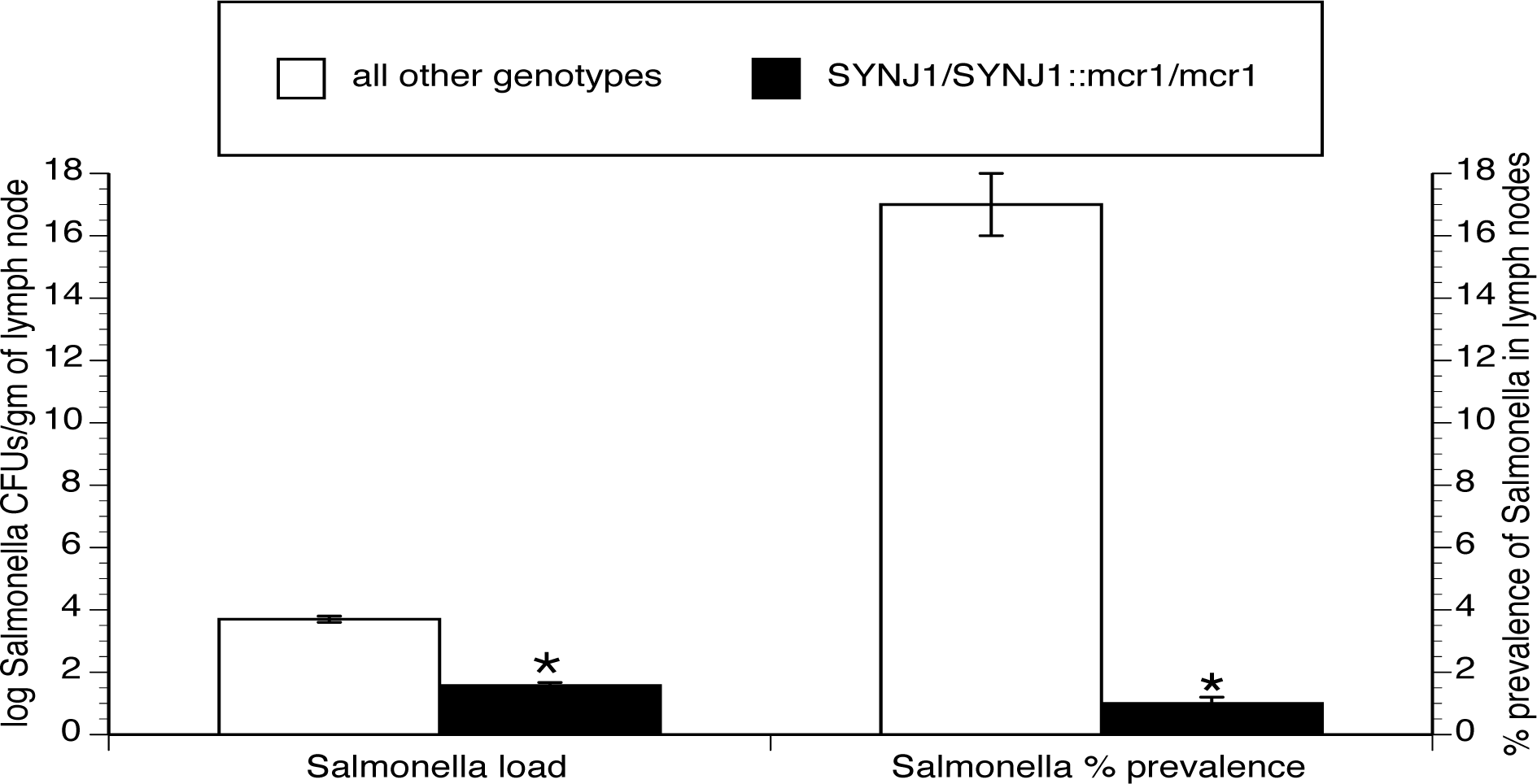
Assessment of lymph node infiltration by *Salmonella* in heifers of various genotypes from a previous study (Feye et al., 2016). *Salmonella* load and prevalence data from the previous study were individually segregated based on the *SYNJ1* and *mcr1* genotypes of each animal. Data represent the mean ± SEM from a total of 400 animals. \**P* < 0.05 versus all other genotypes.

### 3.7. Assessment of in vivo susceptibilities to E. coli O157:H7 colonization across the SYNJ1 and mcr1 genotypes

Calves experimentally infected with *S*. Anatum and Montevideo were also infected with 10^10^ CFUs/kg of *E. coli* O157:H7. On day 14 post-challenge, recto-anal junction scrapings were collected and assessed for the presence of *E. coli* O157H7. As shown in Fig. 7, the load and prevalence of *E. coli* O157:H7 were significantly lower in *SYNJ1/SNYJ1::mcr1/mcr1* cattle.

**Fig. 7.**
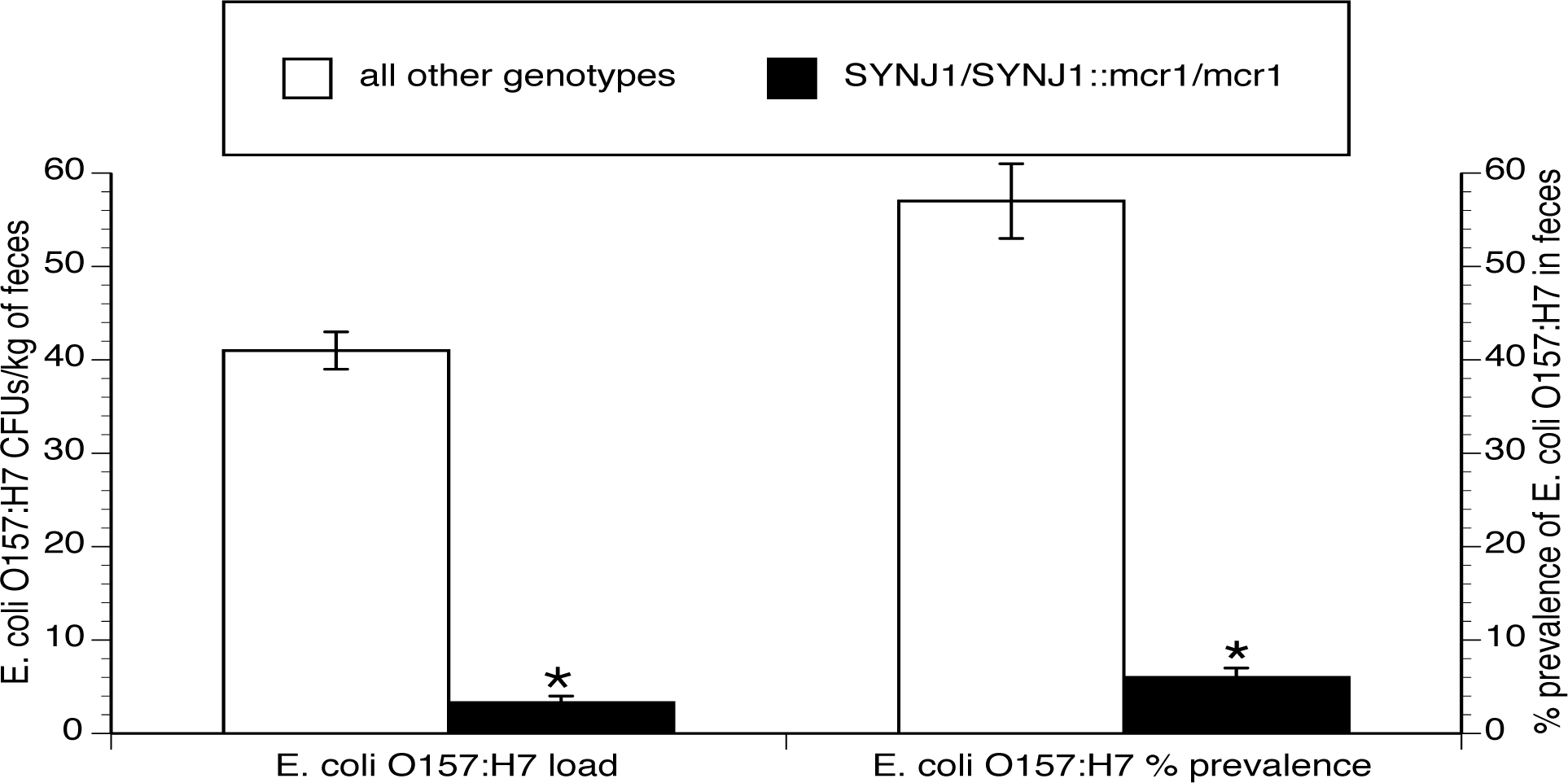
Assessment of recto-anal junction colonization of *E. coli* O157:H7 in *SYNJ1/ SYNJ1::mcr1/mcr1* Cattle. Calves were experimentally infected and recto-anal junction scrapings were recovered and *E. coli* O157:H7 were enumerated in each sample. Data are presented, for scaling purposes, as *E. coli* O157:H7 CFUs/ kg of feces. n=12 for the *SYNJ1/ SYNJ1::mcr1/mcr1* and n=16 for the other eight genotypes (*i.e*., two calves per genotype). \**P* < 0.05 versus all other genotypes.

### 3.8. Small-scale Assessment of synj1 and SYNJ1 Genotypes in Various Cattle Breeds

As shown in Fig. 8, the *SYNJ1/SNYJ1* genotype was present in the following *mcr1/mcr1* breeds: Akaushi, Barzona, Braunvieh, Hereford, Piedmontese, Red and White Holsteins, Red Angus, Red Poll, Shorthorn, Red Simmental, and Tarentaise. In these breeds, the prevalence of the *SYNJ1/SNYJ1* genotype was highly variable but the penetrance of the effect (as determined by *ex vivo* and/or *in vivo* studies) was near 100% in all breeds. For black breeds, the prevalence of the *SYNJ1/SNYJ1* genotype was also highly variable but the penetrance was near zero.

**Figure 8.**
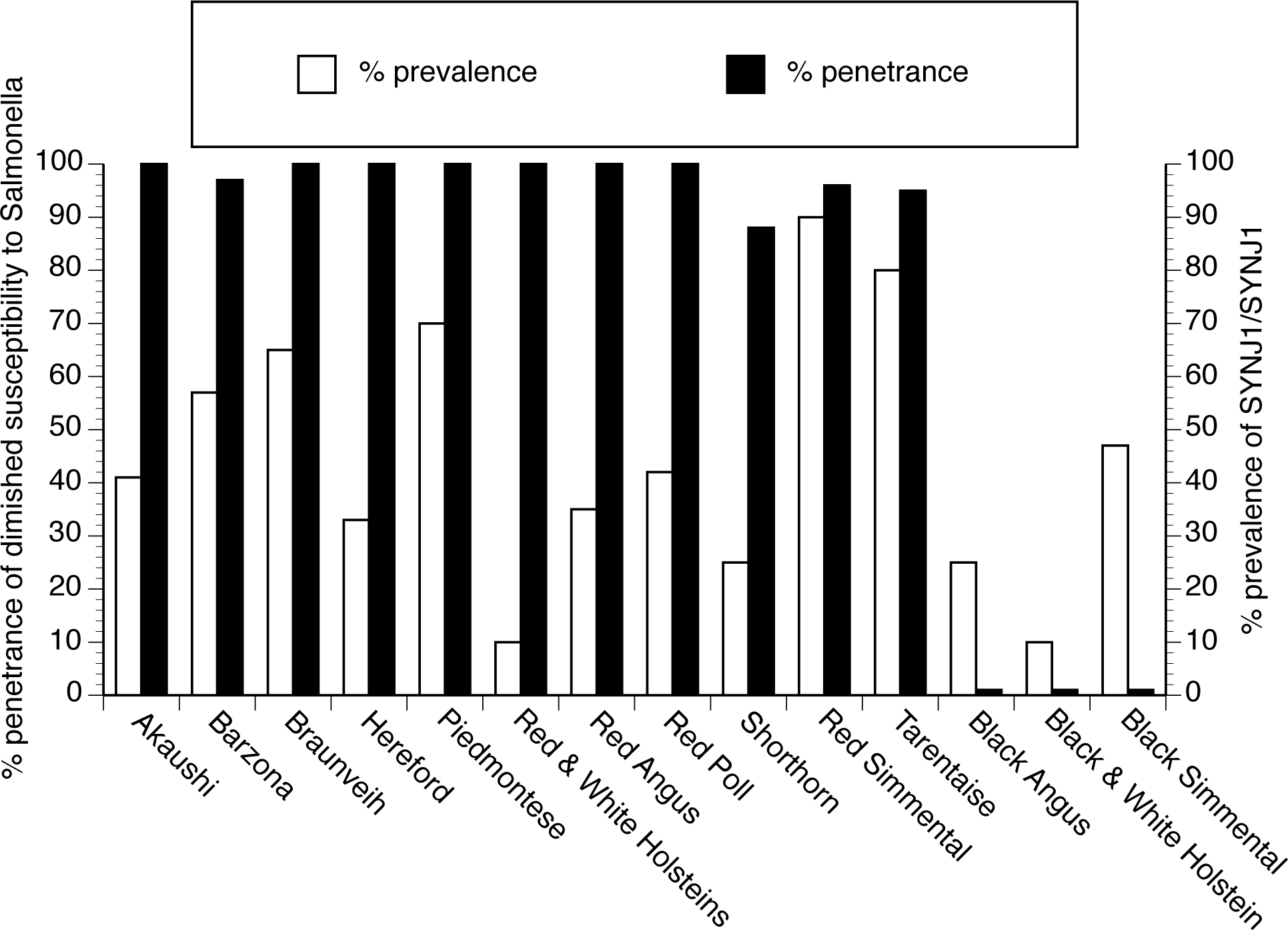
Small-scale assessment of *SYNJ1/SYNJ1* Genotypes and *Salmonella* susceptibility phenotypes in various cattle breeds. Blood was obtained from the following *mcr1/mcr1* breeds: Akaushi (n=24), Barzona (n=30), Braunvieh (n=100), Hereford (n=3), Piedmontese (n=35), Red and White Holsteins (n=20), Red Angus (n=160), Red Poll (n=40), Shorthorn (n=4), Red Simmental (n=20), and Tarentaise (n=40). Blood was also collected from the following *MCR1* breeds: Black and White Holsteins (n=250), Black Angus (n=250), and Black Simmental (n=5). The *ex vivo* invasion and/or *in vivo* susceptibility experiments were performed using the blood samples in order to determine the penetrance of the diminished susceptibility relative to the presence of the *SYNJ1/SYNJ1* genotype. Diminished susceptibility to salmonellosis was ascribed to samples in which the *ex vivo* invasion and survival of the test strain of *Salmonella* (SL1344) was indistinct from that observed for the non-invasive *Salmonella* strain BJ68, or when an animal did not require euthanasia in the *in vivo S*. Newport challenge studies presented in Fig. 4.

### 3.9. Small-scale Assessment of Diminished Susceptibility to an Array of Salmonella serotypes

Since our initial *ex vivo* studies and our *in vivo* studies used just four serotypes, additional studies were performed in order to assess the ability of an array of *Salmonella* serotypes to infect leukocytes from cattle being the *SYNJ1/SNYJ1::mcr1/mcr1* genotype. The *ex vivo* assay, depicted in Fig. 1, was used with a pool of bovine-associated *Salmonella* serotypes incubated with leukocytes from 50 calves bearing the *SYNJ1/SNYJ1::mcr1/mcr1* genotype. Data from each animal was compared to the invasion and survival of SL1344 for that animal. As shown in Fig. 9, about 50% of the samples exhibited enhanced susceptibility to infection when compared to that observed for SL1344. SNP analysis, comparing the 50k SNP profiles of the subset with enhanced susceptibility and reduced susceptibility to the pool of serotypes, revealed that three SNPs underlie this difference. The identities of the SNPs will not be revealed herein and are being investigated further. The “least” susceptibility phenotype was observed at varying frequencies in the breeds shown in Fig. 8.

**Figure 9.**
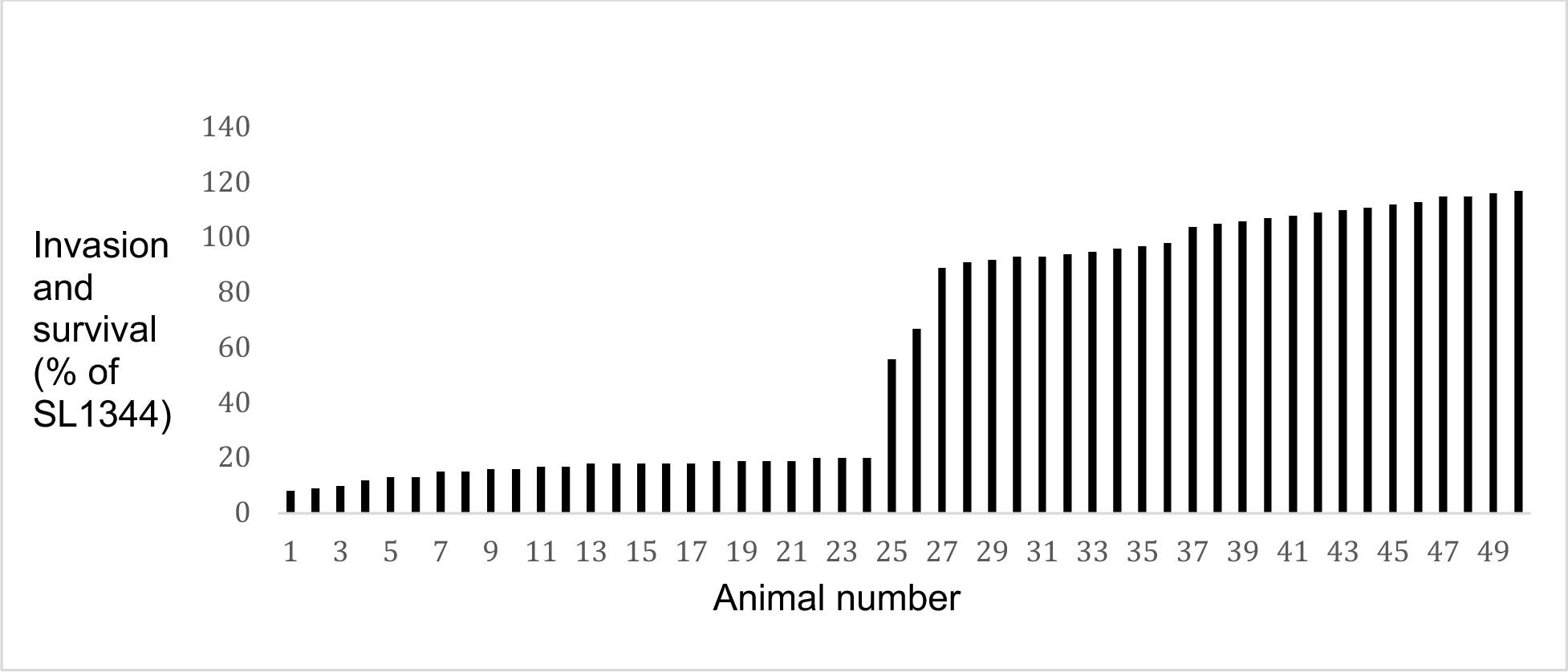
*Ex vivo* studies identify differential susceptibilities to 100 different *Salmonella* strains in *SYNJ1/SYNJ1::mcr1/mcr1* cattle. Leukocytes from 50 different *SYNJ1/SYNJ1::mcr1/mcr1* cattle were incubated with a pool of 100 different strains of *Salmonella* encompassing over 70 serotypes and *Salmonella* were recovered as described in experiments presented in Fig.1. Data were compared, on an individual animal basis, to that observed for the laboratory strain SL1344 that was used as the standard strain in experiments presented in Fig.1.

## 4. Discussion

*Salmonella* and *E. coli* O157:H7 are significant problems for the beef industry, representing two of the most important food safety hazards while *Salmonella* is additionally an animal health issue. Both of these microbes can be shed in feces and *Salmonella* is also present in lymph nodes that contaminate ground beef. Therefore, identifying novel interventions for both pathogens is needed especially considering that lymph node infiltration problem is difficult to mitigate since the lymph nodes cannot be decontaminated (Brichta-Harhay et al., 2008).

In this study, diminished susceptibilities to both pathogens were noted in a very specific subset of cattle that involved a SNP and a coat color genotype. This diminished susceptibility to *Salmonella* was identified using an *ex vivo* assay (Fig. 1) and this “resistance” phenotype was associated with a genotype (Fig. 2) and diminished expression of a bovine protein (Fig. 3) exploited by *Salmonella* during the infection process (Schleker et al., 2012b). The diminished susceptibility was noted for clinical salmonellosis (Fig. 4) and lymph node infiltration (Figs. 5 and 6). The effect also extended to diminished colonization of *E. coli* O157:H7 (Fig. 7).

The basis for the effect revolves around the *SYNJ1/SYNJ1* genotype in which synaptojanin is poorly expressed in bovine cells, and thus it is not an absolute requirement for physiologic functions in cattle. Since *Salmonella* needs to exploit synaptojanin during the infection process, it appears that the paucity of this protein restricts the invasion of the pathogen. However, this paucity alone is insufficient since the *mcr1/mcr1* genotype is also required for “resistance”. That is, the absence of the melanocortin 1 receptor is co-required. The melanocortin 1 receptor has other functions outside of pigmentation, including the beta-defensin and opioid pathways (Leoni et al. 2010). Reducing these other functions may contribute to the elimination of *Salmonella* and the closely related pathogen *E. coli* O157:H7 in *SYNJ1/SYNJ1* cattle, and this reduction is synergistic with the reduction in synaptojanin.

We also identified a few *SYNJ1/SYNJ1* melanotic cattle exhibiting the resistance. These cattle possibly have other SNP(s) in the *Salmonella* interactome, and these other genetic elements may synergize with the diminished expression of synaptojanin.

It is of note that the *SYNJ1/SYNJ1::mcr1/mcr1* genotype does not confer diminished susceptibility to all *Salmonella* serotypes. As shown in Fig. 9, other SNPs are required to extend the diminished susceptibility beyond the four serotypes assessed in Figs. 1-8.

The results of this study identify susceptible and resistant breeds of cattle, however, the molecular mechanisms resulting in these phenotypes has not yet been fully elucidated. It appears that a host protein exploited by *Salmonella* during the infection process (Schleker et al., 2012a) is minimally expressed in cattle with diminished susceptibility. It is also possible that the proteins exploited by *Salmonella* during the infection process are hyper-expressed in the cattle with elevated susceptibility, *e.g*., the melanotic cattle. Ongoing studies will address these hypotheses.

## 5. Conclusions

In summary, this study reveals differing susceptibilities to *Salmonella* infection and *E. coli* O157:H7 colonization in cattle. While we did not examine all breeds of cattle (including those from the *Bos indicus*) and we did not examine haplotypes in the “resistant” cattle, we found several breeds with a significant prevalence of the desired genotype. This work will provide the basis for identifying traits that could possibly be incorporated into cattle, in order to minimize the colonization of *Salmonella* and *E. coli* O157:H7 in the intestinal tracts of a major protein source. PSR Genetics LLC holds a U.S. patent on testing for the *SYNJ1/SNYJ1* genotype (patent number 9,049,848) while the three complementary SNPs are under further investigation.

## Acknowledgments

The authors thank Dr. Ryan Saltzman for assistance in sample collection.

## Funding and Conflict of Interest

This study was funded by PSR Genetics LLC which included a rental agreement for laboratory space at Iowa State University.

